# Comparison of visceral adipose tissue DNA methylation and gene expression profiles in female adolescents with obesity

**DOI:** 10.1101/728832

**Authors:** Matthew D. Barberio, Evan P. Nadler, Samantha Sevilla, Rosemary Lu, Brennan Harmon, Monica J. Hubal

**Affiliations:** Center for Genetic Medicine Research, Children’s Research Institute, Children’s National Medical Center, Washington, DC, United States of America; Sheikh Zayed Institute for Pediatric Surgical Innovation, Children’s Research Institute, Children’s National Medical Center, Washington, DC, United States of America; Division of Pediatric Surgery; Children’s National Medical Center, Washington, DC, United States of America; Department of Integrative Systems Biology, School of Medicine, George Washington University, Washington, DC, United States of America; Department of Kinesiology, Indiana University Purdue University Indianapolis, Indianapolis, IN, United States of America

**Keywords:** Obesity, Adipose Tissue, Epigenetics, Gene Expression

## Abstract

**Background:** Epigenetic changes in visceral adipose tissue (VAT) with obesity and their effects on gene expression are poorly understood, especially during emergent obesity in youth. The current study tested the hypothesis that methylation and gene expression profiles of key growth factor and inflammatory pathways such as PI3K/AKT signaling are altered in VAT from obese compared to non-obese youth.

**Methods:** VAT samples from adolescent females grouped as Lean (L; n=15; age=15±3 yrs, BMI=21.9±3.0 kg/m^2^) or Obese (Ob; n=15, age=16±2 yrs, BMI=45.8±9.8 kg/m^2^) were collected. Global methylation (n=20) and gene expression (N=30) patterns were profiled via microarray and interrogated for differences between groups by ANCOVA (p<0.05), followed by biological pathway analysis.

**Results:** Overlapping differences in methylation and gene expression in 317 genes were found in VAT from obese compared to lean groups. PI3K/AKT Signaling (p=1.83×10^−6^; 10/121 molecules in dataset/pathway) was significantly overrepresented in Ob VAT according to pathway analysis. mRNA upregulations in the PI3K/AKT Signaling Pathway genes *TFAM* (p=0.03; Fold change=1.8) and *PPP2R5C* (p=0.03, FC=2.6) were confirmed via qRT-PCR.

**Conclusion:** Our analyses show obesity-related differences in DNA methylation and gene expression in visceral adipose tissue of adolescent females. Specifically, we identified methylation site/gene expression pairs differentially regulated and mapped these differences to PI3K/AKT signaling, suggesting that PI3K/AKT signaling pathway dysfunction in obesity may be driven in part by obesity-related changes in DNA methylation.

## Introduction

Obesity is a chronic condition characterized by the accumulation of adipose tissue, which plays a critical role in metabolic dysfunction (1). Regionalized accumulation of visceral adipose tissue (VAT) has been strongly linked to the incident development of cardiovascular disease and cardiometabolic diseases such as Type 2 diabetes mellitus (T2DM) and stroke (2, 3). While adipose tissue is recognized as an important endocrine organ responsible for the secretion of multiple pro-inflammatory cytokines and adipokines, the molecular adaptations of adipose tissue to obesity are less clearly understood (4, 5), as are the specific molecular mechanisms driving obesity co-morbidities like insulin resistance.

Heritable and lifestyle (e.g. diet and physical activity) factors play crucial roles in the development of obesity, resulting in a complex pathogenesis of obesity and its co-morbidities (6). Epigenetic regulation represents the intersection between genetics and the obesogenic environment and an increased interest in the role of epigenetics in the development of obesity has developed over the past decade (7). DNA methylation is a key type of epigenetic modification that has received attention because of its tissue-specificity and responsiveness to environmental changes (1). Methylation mainly occurs at the 5’ position of cytosine residues occurring in CG dinucleotide (CpG sites), which are unevenly distributed in the genome. The interest of CpG methylation in complex disease stems from its typical role as a suppressor of gene expression through the methylation of gene promoters (1).

Current evidence suggests DNA methylation is modified across multiple tissues in patients with obesity and obesity co-morbidities (8, 9). Barres et al. (10) demonstrated in skeletal muscle that obesity resulted in genome-wide hyper-methylation of promoter regions that was reversed by weight loss following bariatric surgery. Studies analyzing subcutaneous adipose tissue (SAT) have demonstrated a dynamic methylome in response to acute and chronic bouts of exercise (11), and have linked methylation levels to clinical measures such as blood glucose homeostasis (12). In VAT, over 8,000 CpG sites were found to be differentially-methylated in individuals with obesity and metabolic syndrome, as compared to individuals with obesity but without metabolic syndrome (13). While these, and other, investigations provide insight into epigenetic changes with obesity, these data do not give us a comprehensive pattern of differential methylation with obesity. Data are particularly lacking that couple methylation data with gene expression data in the same samples, which would provide powerful insight into the functional consequences of changes in DNA methylation (14, 15).

Studies addressing adipose tissue global gene expression profiles and global DNA methylation in obesity have been limited to mainly adult cohorts (11–13, 16, 17). In this investigation, we utilize adolescent cohorts with vastly different amounts of VAT, allowing us to identify genomic and epigenetic modifications driven by obesity during this period of dynamic growth and maturation. Understanding how the obesogenic environment affects both DNA methylation and gene expression in VAT tissue with a global unbiased approach can help identify new pathways and molecular targets that can be further tested for their role in tissue dysfunction, as well as their potential targets for therapeutics and diagnostics. We tested the hypothesis that key growth and inflammatory pathways would be epigenetically- and transcriptionally-altered with obesity in adolescent females, as compared to lean age-matched controls.

## Materials and Methods

### Subjects

Adolescent (age 12-19) females grouped as either Lean (L; body mass index (BMI) <25; n=15) or Obese (Ob, BMI >35; n=15) were recruited to participate in this study. Three races (African-American, Caucasian and Hispanic) were represented in equal numbers (5 L; 5 Ob each). Lean patients were recruited from non-bariatric abdominal surgeries (appendectomies and cholecystectomies), while subjects with obesity were recruited prior to bariatric surgery at Children’s National Medical Center. Known clinical diagnoses and medications at the time of surgery are listed below per group – N=1 per diagnosis/medication unless otherwise noted. Lean cohort: sickle cell anemia, localized peritonitis, ulcerative colitis, Crohn’s disease, acid reflux, Remicade (n=2), Albuterol, and prednisone. Obese cohort: asthma (n=4), sleep apnea (n=3), insulin resistance, hypertension, polycystic ovary syndrome, appendicitis, cholecystitis, sickle cell trait, pseudotumor cerebri, hypothyroidism, albuterol, prednisone, Lisinopril, levothyroxine, and Diamox. Bariatric surgery patients had completed a protein-sparing modified fast (~1000 kcal/day; 50-60 g protein) for two weeks prior to their date of surgery. All operations took place after a minimum 12-hour overnight fasting per standard surgical practices. Patients provided assent and legal guardians signed written informed consents as approved by the Children’s National Medical Center Institutional Review Board. Subject characteristics are listed in Table 1.

**Table 1.** Participant Characteristics

| Subjects | Gene Expression Cohort |  |  |  |  |
| --- | --- | --- | --- | --- | --- |
|  | n | Age (yr) | Height (cm) | Weight (kg)* | BMI * |
| **Lean** | 15 | 15 ± 2 | 161 ± 6 | 56.3 ± 11.1 | 21.6 ± 2.7 |
| African American | 5 | 16 ± 3 | 162 ± 8 | 60.3 ± 14.2 | 22.4 ± 3.7 |
| Caucasian | 5 | 16 ± 1 | 162 ± 7 | 55.6 ± 9.3 | 21.2 ± 2.0 |
| Hispanic | 5 | 13 ± 2 | 158 ± 5 | 53.3 ± 9.3 | 21.1 ± 2.0 |
| **Obese** | 15 | 17 ± 2 | 162 ± 8 | 124.4 ± 29 | 47.0 ± 9.7 |
| African American | 5 | 16 ± 2 | 165 ± 7 | 140.2 ± 20.6 | 51.7 ± 9.8 |
| Caucasian | 5 | 16 ± 2 | 163 ± 10 | 111.5 ± 37.7 | 41.5 ± 9.2 |
| Hispanic | 5 | 13 ± 2 | 158 ± 8 | 121.3 ± 23.6 | 47.8 ± 9.0 |
|  | DNA Methylation Cohort |  |  |  |  |
|  | n | Age (yr) | Height (cm) | Weight (kg)* | BMI * |
| **Lean** | 10 | 15 ± 3 | 162 ± 7 | 57.5 ± 2.9 | 21.9 ± 3.0 |
| African American | 4 | 16 ± 3 | 160 ± 7 | 57.2 ± 14.7 | 21.8 ± 4.0 |
| Caucasian | 2 | 16 ± 1 | 166 ± 9 | 64.5 ± 12.2 | 22.9 ± 1.6 |
| Hispanic | 4 | 13 ± 1 | 159 ± 5 | 54.8 ± 10.1 | 21.4 ± 2.6 |
| **Obese** | 10 | 15 ± 2 | 159 ± 7 | 114.2 ± 28.5 | 44.8 ± 9.8 |
| African American | 4 | 17 ± 3 | 164 ± 7 | 148.8 ± 8.7 | 55.6 ± 5.3 |
| Caucasian | 4 | 16 ± 2 | 159 ± 5 | 95.4 ± 12.6 | 37.9 ± 5.0 |
| Hispanic | 2 | 16 ± 1 | 153 ± 9 | 98.2 ± 3.0 | 42.1 ± 6.1 |

\*Significant difference (p<0.05) between lean and obese groups. #Methylation cohort represents a subcohort of the Gene Expression cohort; no statistical significance for phenotype data between the gene expression and methylation cohorts was found.

### Sample Collection

Visceral adipose (VAT) was collected from the omentum during surgery, immediately frozen in liquid nitrogen and stored at −80° C until further processing.

### DNA Isolation and VAT Global DNA Methylation Analysis

A representative sub-cohort of (n=20; 10 L; 10 Ob) of subjects from the entire study cohort (N=30) was chosen for global DNA methylation analysis based on tissue availability and DNA quality. DNA was isolated from ~50 mg of tissue using the QIAamp DNeasy Tissue Kit (Qiagen Inc.; Germantown, MD). DNA quality and quantity were analyzed using the NanoDrop 8000 UV-Vis spectrophotometer (Thermo Scientific; Waltham, MA). Samples all had 260:280 nm ratios > 1.8. DNA was diluted to 25 ng/μl and a total of 500 ng was bisulfate converted with the EZ DNA Methylation Kit (Zymo Research, Orange, CA), using the alternative incubation methods recommended for the Illumina Infinium Methylation Assay (Illumina, Inc., San Diego, CA; Accession: GSE88940). Bisulfite converted DNA (4 ul) was analyzed in a balanced design using Infinium Human Methylation 450 BeadChip Arrays (Illumina, Inc.) per manufacturer’s protocol. BeadChips were scanned on an Illumina iScan System and data were analyzed with Genome Studio software (Illumina, Inc.).

### RNA Isolation and VAT Global Gene Expression Analysis

RNA was isolated from ~200 mg of VAT (N=30) using the RNeasy Lipid Tissue Mini Kit (Qiagen). RNA quality and quantity were analyzed using a NanoDrop spectrophotometer as described above, with all samples having 260:280 ratios > 1.8. Global VAT gene expression was analyzed using Affymetrix Hu133 Plus 2.0 microarrays (Affymetrix, Santa Clara, CA; Accession: GSE88837). Briefly, extracted RNA was twice amplified using Affymetrix GeneChip 3’ IVT Express per manufacturer protocol. Biotinylated cRNA (30 μg) was hybridized to the microarrays. For resultant data, CEL files were imported into Affymetrix Expression Console and CHP files were generated using the PLIER (Probe Logarithmic Intensity Error) algorithm (Affymetrix). Standard quality control measures for adequate amplifications, thresholds for appropriate scaling factors, and RNA integrity (GAPDH 3’/5’ and HSAC07 3’/5’) were used (18). Samples not meeting quality control standards at any point in the above-described process were reprocessed from original total RNA.

### Microarray Data Analyses

VAT DNA methylation arrays were analyzed in Illumina’s Genome Studio software. Raw β-methylation scores (β=intensity of the methylated allele (M) / (intensity of the unmethylated allele (U) + M) + 100) were generated and checked for quality based on manufacturer suggestions. β-values were converted to M-values (M=log^2^(ß/(1-β))), which represent a statistically valid method for analysis of differential methylation (19), using the R statistical environment (20). Differential methylation between Ob and L was assessed using M-values. For ease of biological interpretation, M-values have been converted back to β-values; mean β-values were calculated as the average of all β-values for each group and are presented as % methylation (% methylation=β-values * 100). The Illumina Infinium Methylation Assay contains two separate assays (Infinium I and Infinium II), which perform differently (21). Thus, average DNA methylation for Infinium I and Infinium II probes were calculated and are presented independently. For DNA methylation analysis, probes containing SNPs were removed for further analysis due to cross-reactivity (22). Methylation microarray data have been archived to the Gene Expression Omnibus (GSE88940).

For gene expression analyses, PLIER data were imported into Partek Genomics Suite (Partek, Inc.; St. Louis, MO) and log_2_ transformed for further analyses. Differential global DNA methylation and gene expression between Ob and L groups were analyzed in Partek via 3-way ANCOVA with age, ethnicity, and body mass index (BMI) covariates. Probes with a p<0.05 were considered significant in both analyses and used for further analyses. Gene lists from global DNA methylation and gene expression analyses were then explored for overlapping genes in the Genome Reference Consortium GRCh38 build. Genes found to be significantly different between groups for both global DNA methylation and gene expression were used for biological interpretation via pathway analysis (Figure 1). It is important to note that we chose a more lenient significance (p<0.05) cutoff at the individual gene/methylation site stage because the likelihood of false positive findings is exponentially lowered both by cross-mapping gene expression to methylation results and the use of downstream pathway analyses (which would weed out any random errors due to its reliance on relatedness of results). Gene expression microarray data have been archived to the Gene Expression Omnibus (GSE88837).

**Figure 1.**
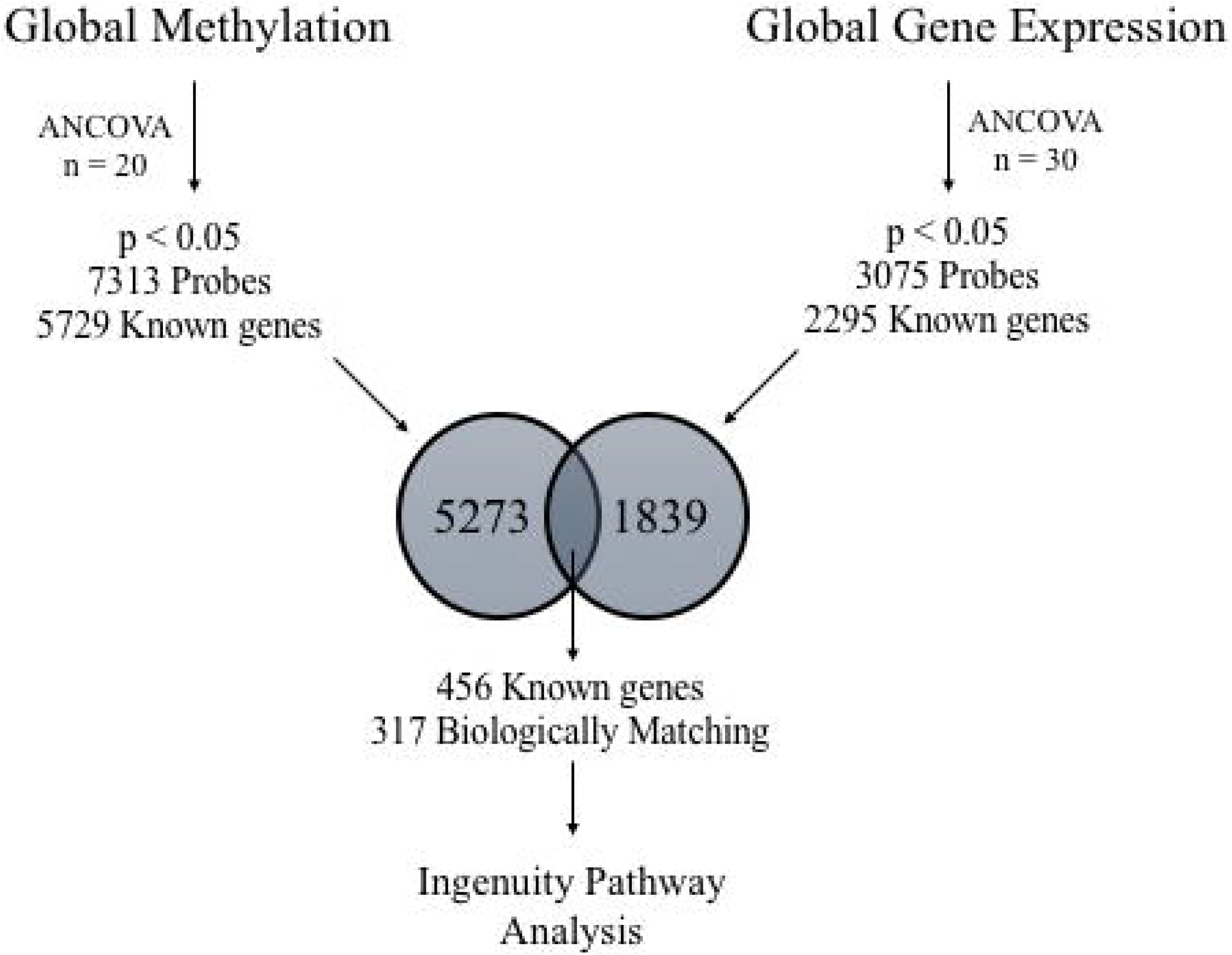
Analysis Work Flow. Genomic DNA and total RNA were extracted from visceral adipose for global DNA methylation analysis (n=20) and global gene expression (N=30), respectively. 3-Way ANCOVA was used to determine (p < 0.05) differential methylation and gene expression. Genes identified in both analyses were further assessed for biological function using Ingenuity Pathway Analysis software.

### Real-Time PCR Validation of Target Genes

Selected microarray results were confirmed via real-time polymerase chain reaction (qPCR) in a larger cohort of 34 (Ob=19, L=15) subjects with available RNA. No statistical differences in demographics were found between this larger cohort and either microarray cohort. RNA (2μg) was reversed-transcribed into cDNA using the SuperScript III Reverse Transcription kit (Invitrogen Corp.; Carlsbad, CA). PCR was performed in triplicate on an Applied Biosystems 7900HT Fast Real-Time PCR System with Taqman Universal PCR Master Mix and commercially available TaqMan human gene expression assays (ThermoFischer Scientific; Waltham, MA) for protein phosphatase 2 regulatory subunit B gamma *(PPP2R5C;* AssayID:Hs00604899_g1) and transcription factor A, mitochondrial *(TFAM;* AssayID:Hs00273372_s1). Assays were performed in accordance with manufacturer instructions: 50°C for 2 min, 95° for 10 min, followed by 40 cycles of 95°C for 15 sec followed by 60°C for 1 min. mRNA content was determined via the comparative C_t_ methodology. Fold changes between Ob and L groups were determined via the 2^−ΔΔCt^ methodology where ΔΔC_t_ = ΔC_t Obese_ – ΔC_t Lean_. Assays were run with a multiplexed endogenous control (18S RNA).

### Biological Pathway Analysis

Ingenuity Pathway Analysis (IPA; Ingenuity, Inc.; Redwood City, CA) was utilized for probe set annotations and to query relationships between genes. For this study, we utilized the canonical pathway analysis tool to identify canonical pathways that were represented in our dataset. Canonical Pathway Analysis utilizes a Right-Handed Tukey’s T-Test to test for overrepresentation of genes/pathways in datasets in comparison to the knowledgebase.

### Other Statistical Analyses

Normality of demographic data was assessed with the Shapiro-Wilk test and visualization of the distribution. If data were non-normally distributed, those data were log_2_ transformed and reassessed for normality. A 2-sample t test was used to assess differences in demographic values between Ob and L cohorts. Differences between sub-cohorts for gene expression and DNA methylation were assessed via 2-sample t test. To maintain independence, subjects represented in DNA methylation cohort were removed from the gene expression cohort for this analysis. Differences in gene expression determined by qPCR were analyzed by via 1-tailed 2-sample t test. Significance was determined *a priori* as p<0.05. Statistical analyses were performed on commercial software (OriginLab Pro 2015; Northampton, MA).

## Results

### Participant Characteristics

Participant characteristics are presented in Table 1. By design, the Ob group had a significantly (p<0.05) higher body mass and BMI (47 ± 10 kg/m^2^ vs. 22 ± 3) than the L group. Groups were similar in age and matched for ethnicity. The sub-cohort of subjects profiled for DNA methylation was representative (no significant differences in age, body mass, or BMI) of the larger gene expression cohort (Table 1). Both cohorts were similar to the qRT-PCR validation cohort (N=34) for all demographic traits.

### Obesity-related Changes in VAT Global Methylation

Globally, 99.98% of the probes on the array were detected and included in analyses. Probes that contain a known single nucleotide polymorphism or did not map to a known gene were removed from analysis, leaving ~300,460 probes in our analysis. ANCOVA detected 7313 probes mapping to 5729 known genes that were differentially methylated (P< 0.05) in Ob as compared to L. Methylation differences between Ob and L VAT ranged from −22% to 26%. Stratification of differentially methylated CpG sites found that 1687 (~33%) sites were located in the gene body, 2149 (~42%) were located in the promoter region (which includes sites within 1500 and 200 bp of the transcription start site; TSS), 189 (4.7%) were located in the 3’UTR, 423 (8.3%) were located in the 1^st^ exon, and 665 (~13.0%) were located in the 5’UTR region. The full list of differentially regulated (p < 0.05) probes from global methylation analysis can be found in Additional Table 1.

### Obesity-related Changes in VAT Global Gene Expression

ANCOVA detected 3,075 probes mapping to 2,295 known genes that were differentially expressed (p < 0.05) in Ob vs L VAT. The full list of differentially regulated (p < 0.05) genes from global gene expression analysis can be found in Additional Table 2. Significant probes were integrated with methylation results to identify overlap to use for Canonical Pathway Analysis within IPA.

### Integration of Global Methylation and Gene Expression Data

Gene lists from methylation and gene expression analyses were combined to identify overlapping genes (Figure 1). In total, 456 genes were identified in both analyses represented by 603 methylation probes and 508 gene expression probes (See Additional Table 3). Probes were then filtered for directionality of change: genes with methylation probes with a negative β_diff_ (indicating hypomethylation in Ob) and gene expression probes with a positive fold change (indicating overexpression in Ob), and vice versa. After filtering, 317 probes remained for biological pathway analysis. The top overlapping genes in Ob as compared to L VAT are presented in Table 2; the full list of overlapping probes for methylation and gene expression is presented in Additional Table 3

**Table 2.**
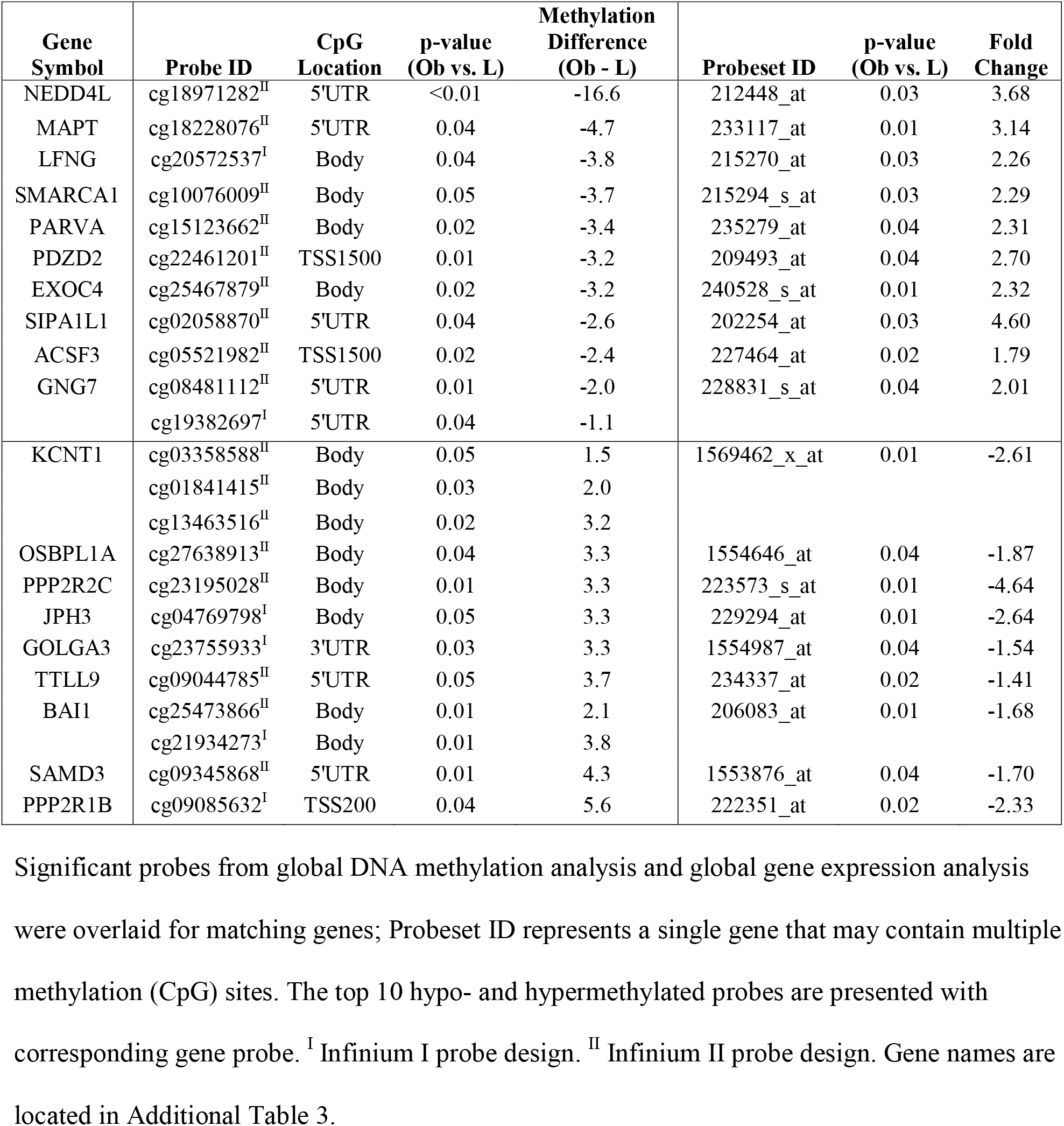
CpG Sites with Highest % Methylation Differences and Corresponding Gene Expression.

### Methylation and Gene Expression Changes in PI3K/AKT Signaling are Altered in Obesity

Differentially regulated genes identified in our overlapping methylation/gene expression analysis were further analyzed using pathway analysis (Figure 2A). Of the 317 probes identified as different between groups in both methylation and expression, 262 were annotated in the IPA databank. We identified PI3K/AKT signaling (p=1.83 x 10^−6^; 11 of 121 genes in the canonical pathway were in the dataset) to be significantly over represented in our gene list (Figure 2B). Methylation and gene expression values for PI3K/AKT Signaling probes are listed in Table 3. To confirm these findings, we confirmed two genes *(PPP2R5C* and *TFAM)* from the PI3K/AKT signaling pathway via traditional qPCR. Statistical analysis indicated a significant increase in VAT expression of *TFAM* (p=0.03, FC=1.8) and *PPP2R5C* (p=0.03, FC=2.6) in the Ob group as compared to the L group (Figure 3).

**Figure 2.**
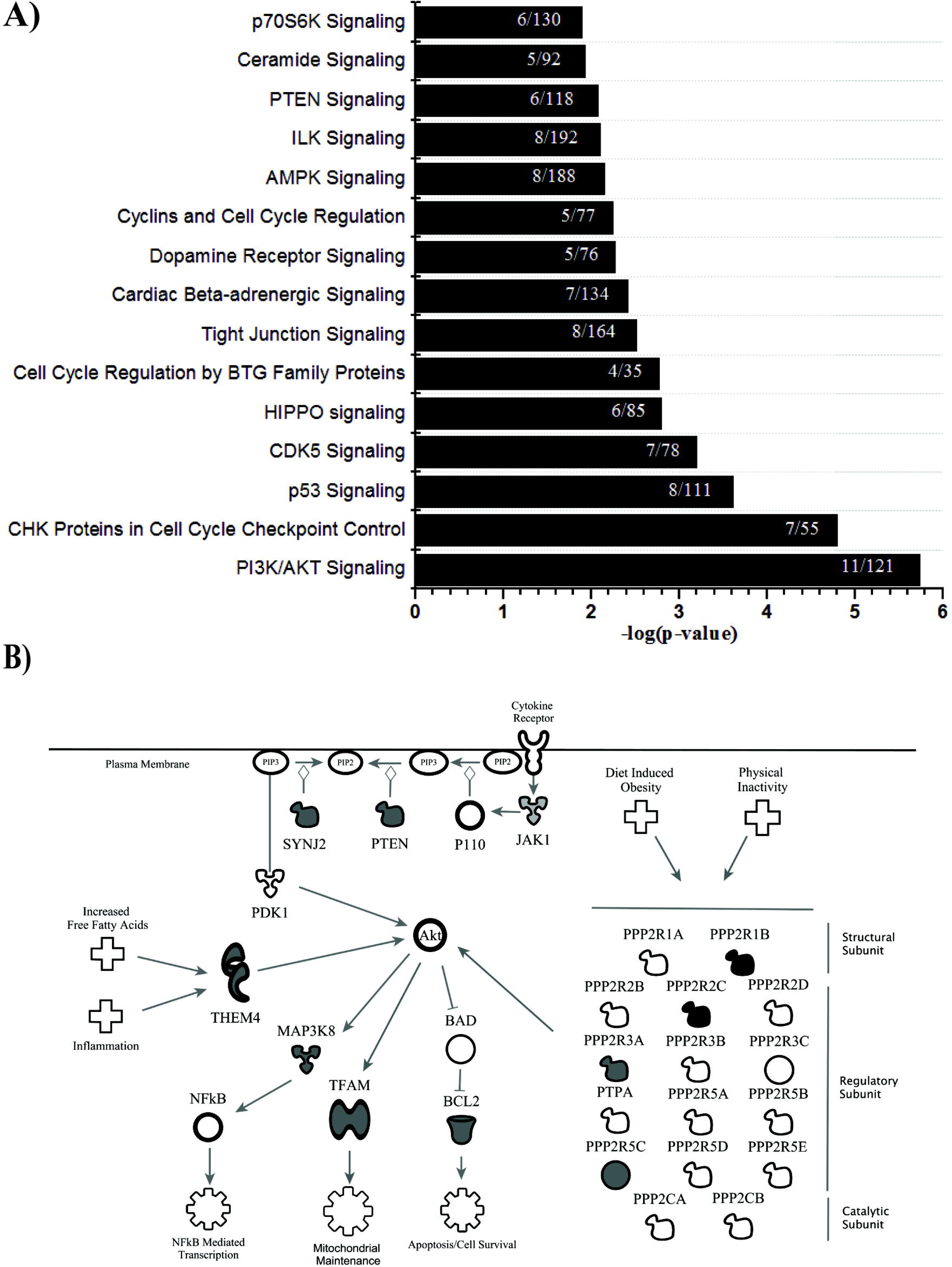
A: Top Canonical Pathways from Ingenuity Pathway Analysis. P-values were determined via Right-Tailed Fisher’s Exact Test and -log transformed. A larger –log(p-value) indicates a lesser likelihood that the grouping of significant genes within the pathway is by random chance. Numbers within bars indicate the ratio of significant genes to total genes within the pathway. **B: Modified IPA PI3K/AKT Signaling with Obesity**. Genes identified in the integrated analysis of global methylation and global gene expression involved in the PI3K/AKT Signaling pathway. Gray shading indicates mRNA upregulation in obesity. Black shading indicates mRNA downregulation in obesity. Corresponding Beta-values, p-values and fold changes are located in Table 3. Figure created using Ingenuity Pathway Analysis.

**Table 3.** PI3K/Akt signaling related genes

| Gene Symbol | Probeset ID | CpG Location | p-value (Ob vs. L) | Methylation Difference (Ob - L) | Probeset ID | p-value (Ob vs. L) | Fold Change |
| --- | --- | --- | --- | --- | --- | --- | --- |
| BCL2 | cg07111678<sup>I</sup> | TSS1500 | 0.04 | -0.33 | 203685_at | 0.04 | 2.5 |
| JAK1 | cg25020373<sup>I</sup> | TSS200 | 0.01 | -0.70 | 240613_at | 0.03 | 2.4 |
| MAP3K8 | cg03807314<sup>II</sup> | TSS200 | 0.04 | -0.19 | 235421_at | 0.04 | 1.9 |
| PPP2R1B | cg09085632<sup>I</sup> | TSS200 | 0.04 | 5.62 | 222351_at | 0.02 | -2.3 |
| PPP2R2C | cg23195028<sup>II</sup> | Body | 0.01 | 3.31 | 223573_s_at | 0.01 | -4.6 |
| PPP2R3A | cg24636867<sup>I</sup> | TSS200 | 0.04 | -0.34 | 207749_s_at | 0.02 | 2.6 |
| PPP2R5C* | cg18997701<sup>II</sup> | Body | 0.02 | -0.29 | 1557718_at | 0.01 | 2.2 |
| PTEN | cg08602305<sup>I</sup> | 5'UTR | 0.03 | -0.31 | 233254_x_at | 0.02 | 6.1 |
| SYNJ2 | cg03682872<sup>I</sup> | TSS200 | 0.03 | -0.35 | 216180_s_at | 0.04 | 1.5 |
| THEM4 | cg03147210<sup>I</sup> | TSS200 | 0.05 | -0.52 | 229253_at | 0.03 | 1.7 |
| TFAM* | cg03859893<sup>II</sup> | TSS200 | 0.05 | -1.02 | 238443_at | 0.05 | 2.5 |

Significant probes from global DNA methylation analysis and global gene expression analysis
were overlaid for matching genes; Probeset ID represents a single gene that may contain multiple
methylation (CpG) sites. IPA Canonical Pathway tool identified PI3K/AKT Signaling as
significantly over represented in our significant gene list. <sup>I</sup> Infinium I probe design. <sup>II</sup> Infinium II
probe design. \* Indicates genes validated via qPCR. Gene names are located in Additional Table

## Discussion

Visceral adipose tissue is a metabolically active endocrine organ that has been linked to the development of obesity and obesity-related comorbidities (23, 24). Understanding the epigenetic and molecular differences in VAT of adolescents with obesity in comparison to lean counterparts may help identify potential therapeutic targets and further inform our understanding of tissue dysfunction in obesity. The use of an adolescent population allows us to identify epigenetic and molecular changes in the earlier stages of obesity, which would be the best targets for prevention/intervention. Using unbiased global methylation and global gene expression analyses, we simultaneously identified epigenetic and functional gene expression changes with obesity involved in the canonical PI3K/AKT signaling pathway.

### Global DNA Methylation Analysis and Gene Expression Analysis

Similar to other published work (8, 9, 13), and using a similar (8) or larger cohort (9), we identified 7313 differentially methylated CpG sites mapping to over 5000 genes. Further, 42% of these CpG sites were located in the promoter region (within 1500 bp of the TSS) while another 33% were located in the gene body of their known genes. We concurrently showed significant differences in 2295 genes through global gene expression analysis. Differential methylation in the promoter region has been demonstrated to be important to gene expression and the metabolic improvement of skeletal muscle following weight loss (10) while gene body methylation is believed to promote alternative slicing or silencing of alternative promoter sites (25). However, there is still a significant amount left to understand about the effect of cell and tissue specific methylation patterns. The degree to which methylation occurs in a particular gene or a particular site is highly variable between cell types and individuals complicating the ability to fully understand the magnitude of differential methylation (Beta difference) of a given site between groups and in different genes. Furthermore, the relationship between specific CpG sites and corresponding mRNA levels remains elusive (15, 26). Our data indicates clear differences in DNA methylation and gene expression patterns in VAT of adolescent females with obesity, indicating a role for epigenetics in driving obesity related changes in gene and tissue function.

### Hyper- and Hypomethylation in PI3K/AKT Signaling Genes

Pathway analysis identified PI3K/AKT signaling as the top canonical pathway represented in our overlapping list of methylation sites/gene expression (Figure 3A and B; individual gene differences detailed in Table 3). Of note in our analysis is that all but one of identified PI3K/AKT genes showed differential methylation in the promoter or gene body, adding to the biological relevance. AKT has a well-defined role in glucose uptake in skeletal muscle and adipose tissue (27). Pathway components included in our gene set are four *(PPP2R1B, PPP2R2C, PPP2R3A*, and *PPP2R5C*) of the 16 genes that encode the heterotrimeric protein phosphatase 2 (PP2A). PP2A is a highly conserved serine/threonine phosphatase involved in the regulation of numerous kinases (28) including AKT. Jun *et al.* (29) showed that PP2A was overexpressed in VAT from rats fed a high fat diet and this resulted in roughly 67% reduction in phosphorylated (i.e. activated) AKT. While we did not measure total PP2A concentrations, we do provide evidence for differential regulation of PP2A genes (−4.6-fold to 2.6-fold) via DNA methylation (−0.34% to 5.6%) in Ob VAT as compared to L VAT. Interestingly, *PPP2R5C* knockout mice have previously been shown to have age-associated obesity (30), though this is likely due to decreased locomotive capacity caused by a heart defect associated with the model. More recently, Cheng *et al.* (31) showed that expression of *PPP2R5C* in the liver correlated with obesity and insulin resistance in patients with obesity with or without diabetes, while knockdown in mice resulted in improved glucose metabolism and alter lipid metabolism. Given the metabolic dysfunction associated with obesity, our finding of a 2.2-fold higher expression of *PPP2R5C* in VAT of Ob as compared to L subjects would further support the conclusion of Cheng *et al.* (31) that *PPP2R5C* is a potential metabolic regulator.

**Figure 3.**
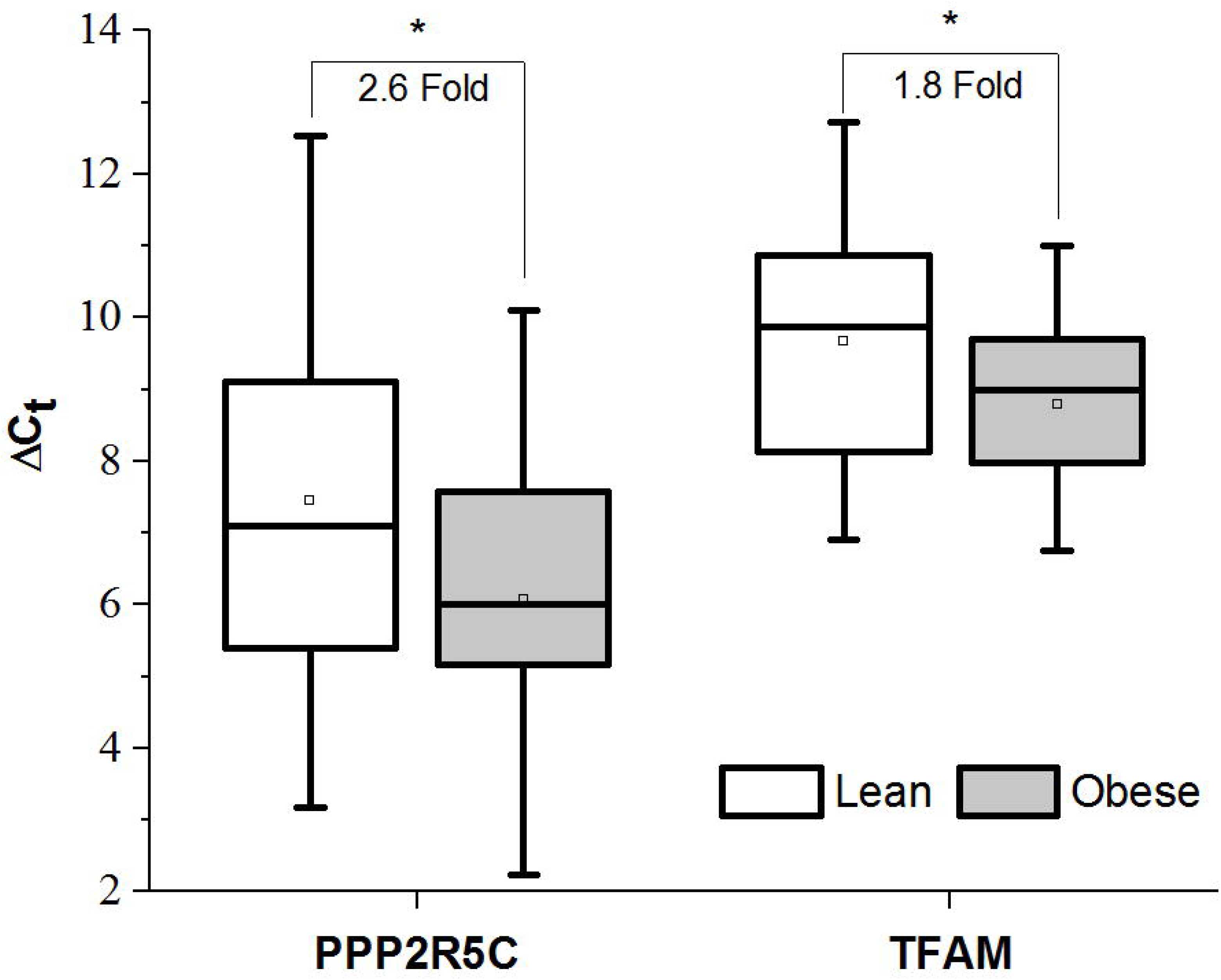
qRT-PCR Confirmation of Select PI3K/Akt Genes. Relative mRNA content of protein phosphatase 2 regulatory subunit B gamma *(PPP2R5C)* and transcription factor A, mitochondrial *(TFAM)* in VAT of lean (white) and persons with obesity (gray) via qRT-PCR. Data are presented as the mean ± SD of the ΔC_t_ (C_t_ target gene – C_t_ endogenous control) for each group. The mean ΔC_t_ for each group is indicated by the open box within each box plot. Lower ΔC_t_ indicates a higher expression. Fold change between groups was calculated as 2^−ΔΔCt^, where ΔΔC_t_ = ΔC_t obesity_ – ΔC_t lean_. * p < 0.05 via 1-tailed two-sample t-test.

We also identified three other PI3K/AKT Signaling related genes previously shown to have significant metabolic and inflammatory effects. Phosphatase and tensin homolog *(PTEN)* was found to be hypomethylated and 6.1-fold higher gene expression in Ob VAT as compared to L. The role of *PTEN* in obesity and metabolic dysfunction remains unclear with studies showing constitutive over-expression resulting in improved energy expenditure (through adipose browning), improved glucose homeostasis, and longer lifespans(32) while adipose-specific deletion (33) also results in improved metabolic parameters. Of further interest are the results of Pal *et al.* (34) that indicate increased risk for obesity and cancer, but a decreased risk for T2DM (mediated through improved insulin sensitivity) in *PTEN* haploinsufficiency. Our analysis also identified mitogen-activated protein kinase kinase kinase 8 *(MAP3K8)*, thioesterase superfamily member 4 *(THEM4)*, and transcription factor A, mitochondrial *(TFAM)* as having differential methylation and gene expression in Ob VAT. *MAP3K8* (also referred to as *TPL2)* was previously shown to be overexpressed in SQ adipose tissue from patients with obesity and this correlated with elevated levels of the inflammatory proteins IL-1β, IL-6, and IL-8 (35). Further exploration of the role of *MAP3K8* in 3T3-L1 and human adipocytes showed activation by inflammatory mediators IL-1β and TNF-α, which affected both lipolysis and ERK signaling (36). *THEM4* (also referred to as *CTMP)* binds to a regulatory domain of AKT, thus preventing its phosphorylation and activation. Here we show VAT from individuals with obesity to have 1.7-fold higher *THEM4* mRNA than lean counterparts. In response to free-fatty acids and inflammatory mediators, THEM4 invokes inhibition of AKT by in immune cells (37) and may participate in the development of impaired insulin resistance in adipocytes (38).

*TFAM* is a key mitochondrial transcription factor important to the activation of mitochondrial transcription. Adipose specific deletion of *TFAM* has been shown to result in protection against obesity and insulin resistance (39) while adiponectin-TFAM-knockout mice were resistant to diet-induced weight gain but suffered from various metabolic abnormalities (40). Obesity-induced mitochondrial dysfunction is a hallmark in multiple tissues and a primary target of pharmaceutical treatment (41, 42). To our knowledge, this is the first study to show evidence of DNA methylation driven changes in *TFAM* mRNA in VAT of individuals with obesity.

While we believe the results and discussion presented in this study are important in furthering the understanding of epigenetic and molecular changes in VAT in obesity, we acknowledge several potential limitations. Only patients with obesity were on a protein-sparing modified fast (as required by the bariatric weight-loss surgical program) and given the environmental responsiveness of DNA methylation and gene expression, we cannot dismiss that potential influence of this diet on our findings, especially since PI3K signaling has been shown to be responsive to such diets (43). However, the response seems to be tissue-(44) and diet-specific (45). Given the biological role of the PI3K/Akt pathways in cellular growth and hypertrophy, we suspect, much like the results of Mercken et al. (37), the diet would have the effect of “normalizing” PI3K/Akt signaling in persons with obesity, which would mean that the differences we found between groups would have been larger if we analyzed samples from pre-diet patients with obesity.

We also did not assess menstrual phase of the patients at the time of the tissue collection (which was logistically impossible for the Lean cohort), though all subjects were post-menarche. Finally, we do not have protein data showing the downstream effects on the reported pathways, as the primary goal of the project was to describe epigenetic changes in obese VAT and how these effects carry over into functional transcriptional changes. Given the invasive nature of VAT collection, we had limited amounts of tissue to work with (especially in Lean controls) to cover both methylome and transcriptome changes, so we did not have extensive tissue remaining for protein studies. Now that we have identified key pathways modified by obesity, future mechanistic studies can address complex protein level changes using a systems biology model.

## CONCLUSION

VAT has long been recognized as a metabolically active and an endocrine-like tissue that releases inflammatory proteins, cytokines, and adipokines. Using unbiased global molecular profiling technologies, we identified obesity-related gene expression changes paired with changes in DNA methylation, noting coordinated changes in the PI3K/AKT signaling pathway, suggesting that PI3K/AKT signaling pathway dysfunction in obesity may be driven in part by DNA methylation as a result of obesity. Further elucidation of the role of DNA methylation in the pathogenesis of obesity-mediated diseases may provide insight into potential therapeutic targets or treatment strategies, especially during early disease development.

## Declarations

Ethics approval and consent to participate: Subjects provided written assent to participation and their legal guardians signed a written informed consent as approved by Children’s National Health System Institutional Review Board (Protocol ID: Pro00000358).

## Consent for publication

Not Applicable

## Availability of data and materials

Gene expression microarray data have been archived to the Gene Expression Omnibus (GSE88837). Methylation microarray data have been archived to the Gene Expression Omnibus (GSE88940). Other data, to the extent reasonable for the protection of Private Health Information, may be available upon request.

## Sources of Funding

This project was supported by Award Number UL1TR000075 (MJH) from the NIH National Center for Advancing Translational Sciences and T32AR065993 (MDB) from the National Institute of arthritis and Musculoskeletal and Skin Diseases. Its contents are solely the responsibility of the authors and do not necessarily represent the official views of the National Center for Advancing Translational Sciences, National Institute of Arthritis and Musculoskeletal and Skin Disease, or the National Institutes of Health. The funders had no role in study design, data collection, and Competing interests: The authors have no competing interest to declare.

## Authors contributions

All authors have agreed to be accountable for all aspects of this work including the accuracy and integrity of all data and information. All authors have given final approval of this manuscript for publication. MDB, EPN, MJH were involved in all aspects of this and and manuscript. SS and RL were involved with sample collection, sample processing. SS, RL, and BH were all involved in the production and analysis of data along with production of the final manuscript.

## Acknowledgements

The authors would like to acknowledge Jelena Pervanovic Ph.D. for assistance with methylation data analyses and Margaret Morrison for her assistance with the project.

---

**Additional Table 1. Differentially regulated DNA methylation probes from global DNA methylation analysis.**

**Additional Table 2. Differentially regulated genes from global gene expression analysis.**

**Additional Table 3. All overlapping probes/genes from DNA methylation and global gene expression analysis.**

## Notes

https://www.ncbi.nlm.nih.gov/geo/query/acc.cgi?acc=GSE88940

https://www.ncbi.nlm.nih.gov/geo/query/acc.cgi?acc=GSE88837

